# Differences in Gut Microbiota Assembly Alter Its Ability to Metabolize Dietary Polysaccharides and Resist *Clostridioides difficile* Colonization

**DOI:** 10.1101/2022.05.20.492827

**Authors:** Matthew K. Schnizlein, Alexandra K. Standke, Mark J. Garmo, Summer J. Edwards, Vincent B. Young

## Abstract

The mammalian gut is home to a vibrant community of microbes. As the gut microbiota has evolved, its members have formed a complex yet stable relationships that prevent non-indigenous microorganisms, such as *Clostridioides difficile*, from establishing within the gut. Using a bioreactor model of the gut, we characterize how variation in microbial community assembly changes its ability to resist *C. difficile*. We established diluted microbial communities from healthy human stool in a bioreactor gut model and subsequently challenged them with vegetative *C. difficile*. 16S rRNA-gene sequencing and selective plating revealed that dilution progressively increases microbiota variability and decreases C. *difficile* colonization resistance. Using Dirichlet Multinomial Mixtures and linear discriminant analysis of effect size, we identified 19 bacterial taxa, including *Bifidobacterium, Bacteroides* and Lachnospiraceae, that associate with more resistant community types. Since these taxa are associated with butyrate production, which is tied to *C. difficile* colonization resistance, we performed another reactor experiment where we increased inulin concentrations prior to *C. difficile* challenge. Diluted communities concurrently lost their ability to produce additional butyrate in response to inulin, as measured by high performance liquid chromatography, and resist *C. difficile* colonization. These data demonstrate that a similar level of microbiota cohesiveness is required to prevent *C. difficile* colonization and metabolize inulin. It also suggests that metabolic activity of butyrate-producing microbes is tied to colonization resistance. Future work can leverage these findings to develop treatments that leverage knowledge of these ecological dynamics to improve efficacy.

**Importance:** The microbes living in the human large intestine helps create an environment that is resistant to organisms that do not normally reside there, such as the pathogen *Clostridioides difficile*. Differences in ways in which microbial communities make an environment their home can change their ability to provide that resistance. To study those differences, we use a model of the intestine that incorporates only microbial variables (i.e. no host is involved). By diluting microbial communities to decrease their complexity, we show that communities lose their ability to resist *C. difficile* at a particular point and, at the same time, their ability to use inulin, a common dietary fiber, in ways that make the environment more toxic to *C. difficile*. These findings will help future researchers dissect the microbial components that create a resistant intestinal environment.

## 1. Introduction

The mammalian gut contains a complex ecosystem with a variety of fungal, bacterial, archaeal and viral organisms that exist in a network of metabolic interactions. These vast arrays of interactions regulate microbial competition and impact the host. External stability, otherwise known as colonization resistance, is a complex phenomenon in which resident taxa prevent the invasion of foreign ones by occupying niches in an environment. For example, probiotic organisms fail to exist long-term in the gut because resident microbes are better able to compete for niche space (1). These interactions have also been observed to prevent colonization of pathogenic organisms such as *Escherichia coli, Salmonella* Typhimurium, and *Clostridioides difficile* (2-5).

In studying colonization resistance, several models, including humans, mice, enteroids and bioreactors, have been used to ascertain characteristics of a resistant environment (5-13). Each of these models supports a unique level of complexity that may consider host-microbe, microbe-microbe, and microbe-environment interactions. Bioreactors have been used extensively to study the microbiota of the gut environment, particularly the dynamics of microbe-mediated colonization resistance. This is due in part to the controlled way nutrients flow in and out of the system. In the context of resistance to *C. difficile*, healthy human stool established in these reactors prevents or limits colonization (14, 15). However, alterations to the resident microbiota can reduce the ecosystem’s ability to do so (15-17).

Since much of colonization resistance revolves around microbial metabolism, host dietary inputs play an important role in modulating this phenotype. As previous studies have demonstrated, both macronutrients (e.g., proteins and polysaccharides) andn cofactors (e.g., vitamins and minerals) can modulate *C. difficile* colonization resistance by affecting the resident microbiota and host immune system (18-20). Much like “macroorganisms” adapt to the food available to them, microbes alter their metabolism to capitalize on the nutrient sources in their surroundings. These metabolic shifts lead to different downstream products with which other organisms in the ecosystem interact.

While previous work has characterized forced effects (e.g., antibiotic use) on microbiota post-establishment, limited work has characterized the stochastic processes of community establishment in the context of *C. difficile* colonization resistance (14, 21, 22). To manipulate the microbiota to treat infections, a fuller grasp of the ecological rules underlying community physiology is needed. Specifically, further work is required to accurately describe how random and specific effects alter community establishment and ability to resist the colonization of non-indigenous microbes. Here we describe how stochastic effects, as induced by community dilution, and directed effects, as induced by supplementation with additional carbohydrates, influence the establishment of a microbial community. We also characterize how the functional effects of this variation alter metabolic output and colonization resistance to *Clostridioides difficile*.

## 2. Results

### 2.1. Dilution of starting inoculum alters establishment dynamics of continuous flow cultures

Using a bioreactor system initially developed by Auchtung et al., we extended studies on how dilution impacts the membership and variability of microbial communities (14, 22). We established communities in bioreactors from serially diluted stool samples taken from a healthy human donor (10^−3^, N=4; 10^−5^, 10^−7^, and 10^−9^, N=6). Following inoculation of the reactors, communities were given one day to equilibrate in static culture before initiating continuous flow. The subsequent 6 days in continuous flow allowed communities time to adjust before testing their external stability with a model invasive bacterium, *C. difficile* (Fig. 1A).

**Figure 1:**
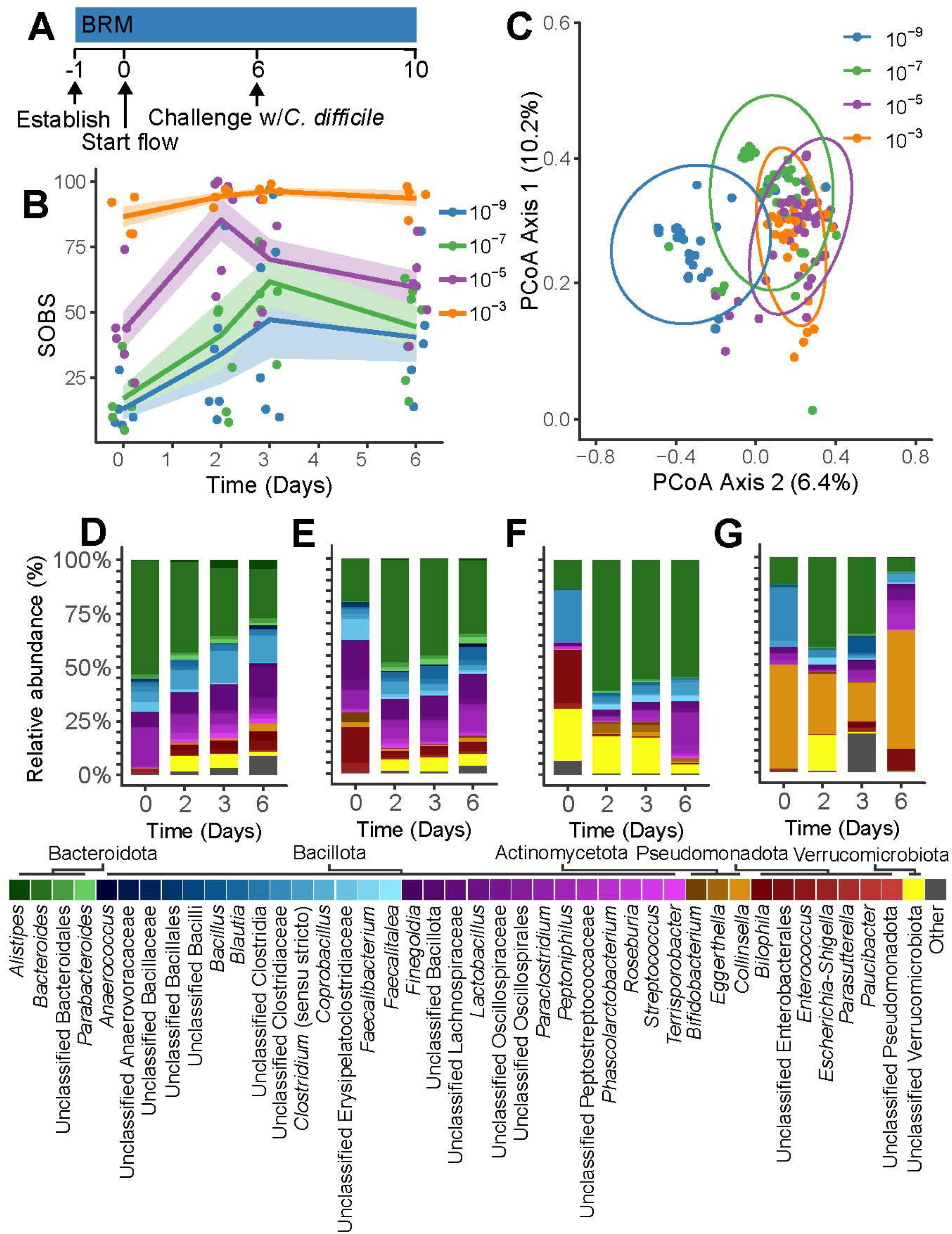
Establishment dynamics of diluted microbial communities. **A)** Experimental timeline of the bioreactor dilution experiment. **B)** Observed species (SOBS) is plotted by dilution from day 0 to the day of *C. difficile* challenge. **C)** Principal coordinate analysis of all timepoints in each community dilution (ellipses represent the 95% confidence interval of a multivariate t-distribution for datapoints in each dilution). **D-G)** Averaged relative abundance at each indicated timepoint for the communities diluted **D)** 10^−3^, **E)** 10^−5^, **F)** 10^−7^, and **G)** 10^−9^. Taxa are color coded and ordered by phylum (Bacteroidota = green, Bacillota = blue/purple, Actinomycetota = orange, Pseudomonadota = red and Verrucomicrobiota = yellow. Other phyla, low abundance taxa and unclassified bacteria are colored as grey).

Dilution increased the variability of communities and lowered the number of taxa that became established. By day 6, 16S rRNA-gene sequencing analysis showed that 93±6 and 60±15 operational taxonomic units (OTUs) became established in those communities diluted 10^−3^ and 10^−5^, respectively (Fig. 1B & Fig. S1A). Communities from these stool dilutions consisted mainly of Bacteroidota and Bacillota (Fig. 1D-E). Reactors established with more diluted fecal inocula had fewer OTUs established by day 6 (i.e., 10^−7^ and 10^−9^ dilutions had 45±20 and 40±23 OTUs, respectively) and also unique proportions of bacterial phyla, with some being dominated by Actinomycetota and others by Pseudomonadota (Fig. 1B & Fig. 1F-G). More diluted inocula established in individualized community structures in each reactor replicate (Fig. S1C-F).

This variability is captured by principal coordinate analysis, which shows that dilution altered the dynamics of each community’s establishment so that they cluster separately (Fig. 1C). Dissimilarity between replicate reactors in each group increased as the dilution factor increased, as measured by Bray-Curtis and Jaccard Dissimilarity Indexes, which capture the abundance and the presence/absence of taxa, respectively (Fig. 2A-B). This variability trend was also observed when comparing multiple timepoints within each individual reactor (Fig. S2A-B). While dilution greatly reduced the initial biomass of microbes, after 6 days of growing in continuous culture, communities had reached similar levels of abundance as measured by qPCR of the 16S rRNA-gene (Fig. S1B).

**Figure 2:**
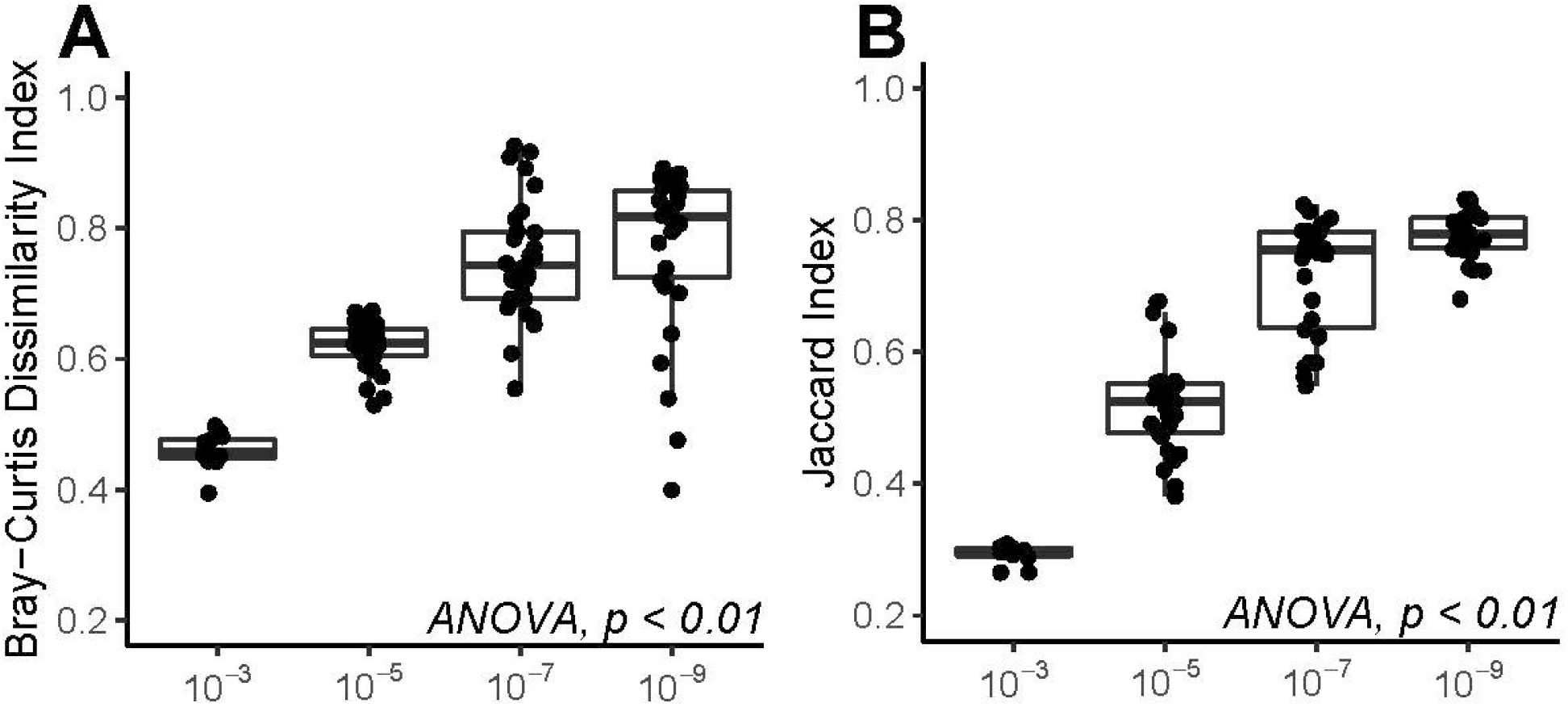
Intra-group community variability by dilution. Intra-group β-diversity was compared across samples for each of the four community dilution groups using the **A)** Bray-Curtis Dissimilarity Index and the **B)** Jaccard Dissimilarity Index (statistical analysis: one-way ANOVA).

### 2.2. Dilution decreases resistance to a model invasive organism

*C. difficile* is a model organism that can generally not invade communities unless they have been perturbed. Since dilution increased community variability, which is a marker of external stability, we hypothesized that this would result in reduced ability to prevent *C. difficile* colonization. 7 days after establishment of bioreactor communities, they were challenged with 10^4^ vegetative *C. difficile* cells. As measured by colony-forming units (CFU), communities possessed varying capabilities to resist colonization with *C. difficile* (Fig. 3A). Within 24 hrs of challenge, 3 of 4 communities diluted 10^−3^ prevented *C. difficile* colonization while communities diluted 10^−9^ showed colonization levels around 10^7^ CFU/mL in all six replicates. The largest intra-group variation in colonization was observed in the 10^−5^, where all six reactors had intermediate levels of colonization, and 10^−7^ dilutions, which had three reactors colonize at 10^7^ and three fully resist. Since *C. difficile* colonizes at 10^7^ CFU/mL when it grows by itself in a reactor (data not shown), our data suggest that reactor communities that reached this level had no colonization resistance. Furthermore, 24 hrs after *C. difficile* challenge, those communities experienced a loss of resident taxa (37±21 on day 6 to 14±4 on day 7; Wilcoxon Rank Sum Test, *p* < 0.01), demonstrating that these microbial communities had both no ability to resist a non-indigenous microbe or remain intact (Fig. S3A-C).

**Figure 3:**
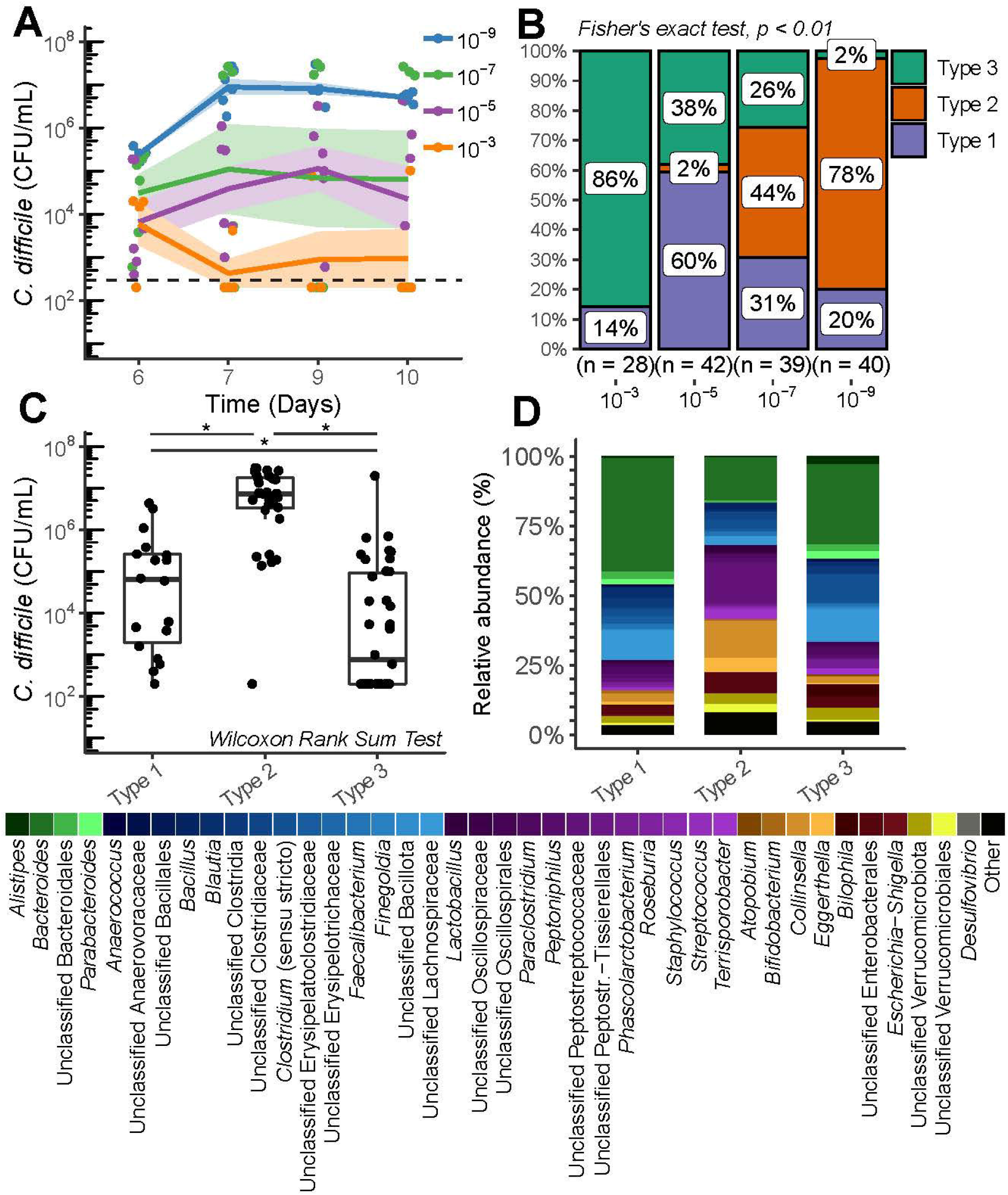
Associations between *C. difficile* colonization and microbiota community type. **A)** *C. difficile* colonization on each day following challenge measured in CFU/mL (dashed line = limit of detection). **B)** All samples were categorized into three unique community types by Dirichlet Multinomial Modelling (DMM). Their relative abundance in each dilution group is plotted with the total number of samples in each dilution listed on the x-axis (statistical analysis: Fisher’s Exact Test). **C)** *C. difficile* abundance in each sample is compared to that samples associated community type (statistical analysis: Wilcoxon Rank Sum Test, * indicates *p* < 0.01). **D)** Average relative abundance of bacterial taxa in each community type (Peptostreptococcales is abbreviated as Peptostr.; Bacteroidota = green, Bacillota = blue/purple, Actinomycetota = orange, Pseudomonadota = red, Verrucomicrobiota = yellow and Desulfovibrio = grey. Other phyla, low abundance taxa and unclassified bacteria are colored as black).

Using Dirichlet Multinomial Modelling (DMM), we identified 3 community types across the established communities, which associated with dilution (Fig. 3B). Of these, type 3 supported significantly lower colonization than enterotypes 1 and 2, with type 1 supporting a middle level of colonization (Fig. 3C). Through LEfSe, we identified 19 taxa associated with the more resistant enterotypes (i.e., enterotypes 1 & 3; Fig. S3D) and five associated with enterotype 2 (Fig. S3E). We noted several commonly associated with metabolic functions known to increase resistance to *C. difficile*, such as short-chain fatty acid (SCFA) production. These taxa included *Bifidobacterium, Bacteroides, Blautia, Faecalibacterium*, Unclassified Lachnospiraceae and *Clostridium* (sensu stricto) (Fig. 3D & Fig. S3D-E).

### 2.3. Diluted communities respond uniquely to a change in carbohydrate concentrations

The dilution experiments above indicated that colonization resistance against *C. difficile* was associated with the presence of taxa that could degrade dietary polysaccharides and produce SCFAs (20, 23). Other work has indicated that SCFAs are able to limit the growth of *C. difficile* (20). Therefore, we characterized how a change to higher carbohydrate concentrations affected the formation of communities and their ability to resist *C. difficile* colonization following a bottleneck event. We chose to increase the availability of the carbohydrate inulin due to its ability to induce the production of SCFAs by the gut microbiota (24-26). We also opted to use this polysaccharide due to the association of microbes with inulin catabolic potential and colonization resistance in the dilution experiment. Using a second fecal donor, we established reactor communities using feces diluted 10^−3^, 10^−4^, 10^−5^ and 10^−6^. Based on our work above we hypothesized that these dilutions would result in communities that might support moderate levels of *C. difficile* colonization. After growing communities for two days on standard BRM, which contains 0.02% inulin, we increased inulin concentrations to 0.2% (Fig. 4A). Despite the narrower range of dilution, communities established in these reactors again differed in the number of OTUs that became established, with less diluted communities having a significantly higher number of resident OTUs (Fig. 4B). While we observed a shift in the Bray-Curtis distance relative to baseline in the community diluted 10^−3^ when comparing days 2 and 5, we did not observe a statistically significant change in the number of OTUs (Fig. 4B & Fig. S4A). There was no shift in ecologic distance in the other dilution groups. Despite the minimal changes in community structure, there was a significant shift in metabolic activity of these communities in terms of carbohydrate metabolism with a drop in acetate and a 3-fold increase in butyrate concentrations (Fig. 4C). There was no change in propionate concentrations (Fig. 4C). Interestingly, butyrate production prior to the shift in inulin (i.e., day 2) was predictive of concentrations afterward (i.e., day 7), with only communities diluted 10^−3^ and 10^−4^ responding to higher inulin (Fig. 4D & Fig. S4C). Despite this variation in metabolic output, pH did not differ across dilutions when measured at day 10 (Fig. S4B). Our data suggest that, in our bioreactor system, a 10-fold inulin increase altered the functional output of the community with minimal effect on its composition, with more diluted communities not responding to that change. Four days after challenging with 10^5^ vegetative *C. difficile* cells, reactors colonized to an average density of 10^6^ CFU/mL *C. difficile* in the 10^−5^ and 10^−6^ groups (Fig. 4E). While all reactors had low initial levels of *C. difficile* colonization, only the communities in the 10^−3^ and 10^−4^ groups were able to ultimately prevent *C. difficile* invasion, suggesting that communities still able to respond to inulin also had the metabolic functions required to mediate colonization resistance (Fig. 4E). While we observed a correlation between butyrate concentrations at day 0 and *C. difficile* colonization at day 4, butyrate did not affect *C. difficile* at the concentrations measured in our reactors in *in vitro* curves at pH 7 (Fig. S4D-F).

**Figure 4:**
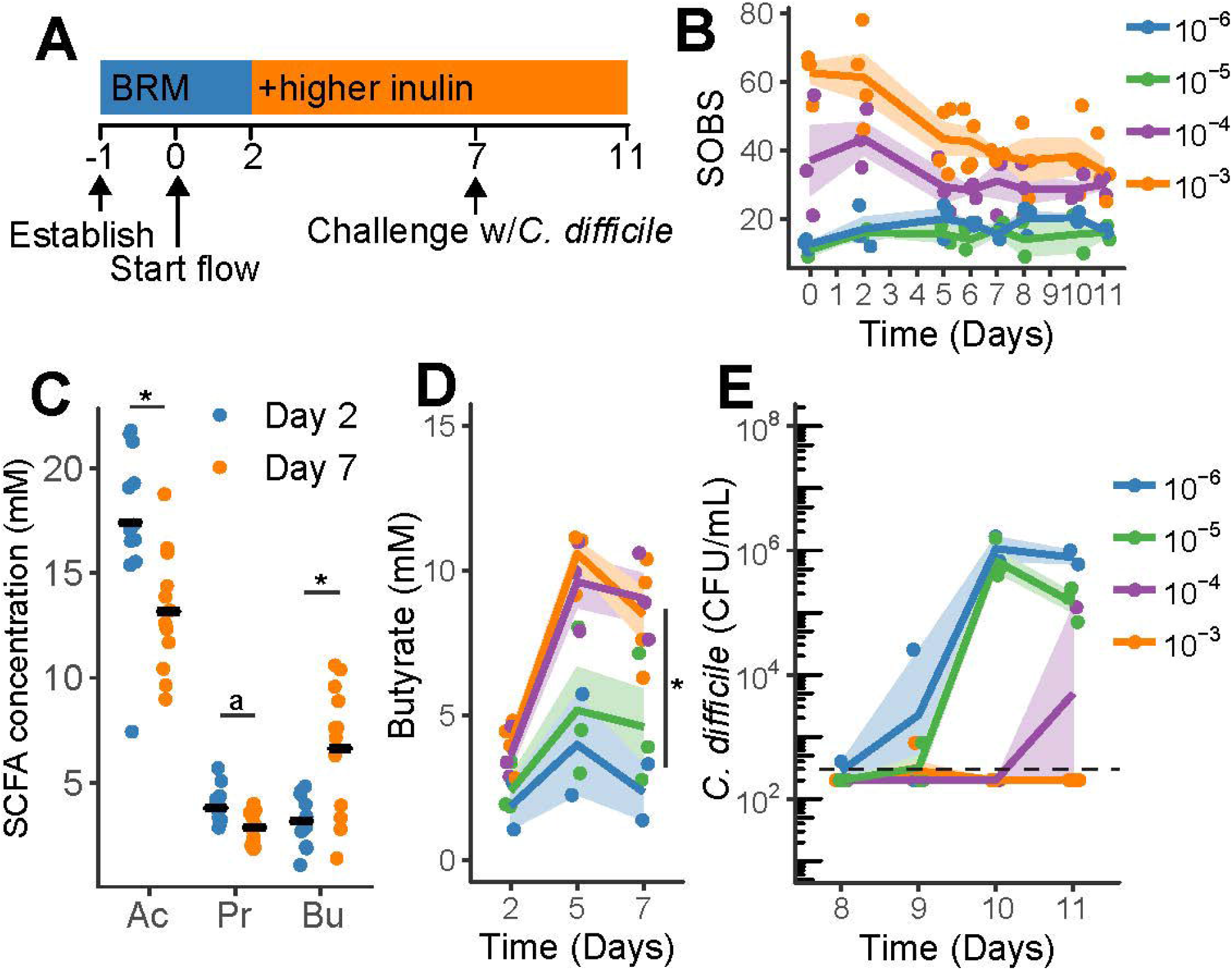
Effect of inulin on microbial community function. **A)** Experimental timeline of the bioreactor carbohydrate experiment. **B)** Observed species (SOBS) over time colored by dilution group (statistical analysis: Kruskal-Wallis Test comparing SOBS at all timepoints with dilution group, *p* < 0.01; Wilcoxon Rank Sum Test comparing OTUs in each group at day 2 and day 5, *p* = not significant). **C)** Short chain fatty acid concentrations in the 0.2% inulin group as measured by HPLC from day 2 (pre-media switch) and day 7 (5 days post-media switch; Ac = acetate, Pr = propionate, Bu = butyrate; statistical analysis: Wilcoxon Rank Sum Test, * indicates *p* < 0.01 and a indicates *p* = 0.033). **D)** Butyrate response 3 and 5 days following the shift to higher inulin, with communities being colored by dilution (statistical analysis: at day 7, 10^−3^ and 10^−4^ compared to 10^−5^ and 10^−6^ using the Wilcoxon Rank Sum Test, * indicates *p* < 0.01). **E)** *C. difficile* colonization in the reactors treated with 0.2% inulin, colored by community dilution (dashed line = limit of detection).

## 3. Discussion

The ability of an ecosystem to resist invasion by a non-indigenous species is tied to the diversity and temporal variability of community membership, an observation noted throughout the animal kingdom (27-30). Whether at the “macro” or “micro” levels of biology, environments provide a given set of nutrients to resident organisms, creating multi-dimensional niches comprised of biotic (e.g., nutrients, predators, etc.) and abiotic (eg., space, gas gradients, etc.) factors (31-36). The diversity and variability of a resident community determines how efficiently niches in the surrounding environment are utilized (36). In our study, we demonstrated that dilution of a community increases the variability of the established community, and those shifts are associated with *C. difficile* colonization resistance. However, this variability is only a marker of underlying metabolic interactions of colonization resistance as evidenced by an increase in resistance when we altered nutrient inputs into that environment.

These metabolic interactions manifest externally through greater competition with invaders as well as internally through limits placed on opportunistic taxa already present in that environment. Thus, we observed dilution having a two-fold disruptive effect on establishing communities, each tied to niche availability. First, while diversity is not strictly a metric of external stability, greater diversity allows for higher numbers of unique interactions, both mutualistic and antagonistic, among community members (27, 37). These interactions increase the likelihood that spaces within a given niche become occupied through stronger cross-feeding interactions between resident members, which limits the invasion of foreign microbes (38-40). By removing rarer taxa, the stochastic nature of dilution weakened these interactions by destabilizing how the remaining microbes co-adapted to their new environment. The resulting changes decreased niche coverage by the resident microbiota (38). The resulting gaps reward microbes like *C. difficile*, which can adapt to use distinct niches depending on what is available in an environment (41). Additionally, networks of mutualistic and antagonistic interactions between microbes increase the likelihood of a microbial community’s long-term survival (42). As observed in the communities that fully colonized with *C. difficile*, invasion by a foreign microbe triggered a collapse in the networks between resident organisms, resulting in their extinction (42).

Second, in addition to removing rarer taxa, dilution decreased the biomass of microbes starting off in each reactor. Since microbial abundance had recovered by the time of *C. difficile* challenge, we do not think that low microbial density played a direct role in the niches available to *C. difficile*. However, low microbial density left large open niches at the outset, which altered the early dynamics of community establishment. Microbes arrived in an environment absent of the competitors that previously had limited their expansion. This founder effect allowed opportunistic taxa within the resident community to take on outsized proportions due to their ability to use surrounding resources more efficiently (43, 44). Since the density of a seeding community regulates how they establish, it is inherently tied to how environmental niches become occupied (45). For example, broad-spectrum antibiotic treatment induces significant gaps in niche coverage by reducing microbial abundance (46, 47). The downstream effects of these perturbations can linger for years, particularly if the event occurs early in the stages of microbial community development (e.g., in human infancy) (48). While perturbations that occur after a community reaches “maturity” have persistent effects, microbial communities tend to be impacted to a lesser extent (49, 50).

In our study, we also investigated the role of carbohydrates and microbial short-chain fatty acid metabolism in revealing the nature of altered community assembly affected by founder effects. While the metabolic nature of colonization resistance is multifactorial, several studies have characterized the role of SCFAs in limiting *C. difficile* establishment in the gut by altering the physiology of both microbe and host (23, 51, 52). While all SCFAs we measured can be products of inulin degradation, butyrate is relevant due to its toxicity to *C. difficile* as well as its ability to limit toxin-associated damage on the colonic epithelium (20, 51). While higher inulin concentrations induced an increase in butyrate, *C. difficile* tolerated those butyrate concentrations as measured by *in vitro* growth curves. Previous work suggests that SCFAs affect bacterial cells in a pH-dependent mechanism, with higher toxicity at lower pH due to the protonated acid form passing more easily through cellular membranes (53, 54). Our *in vitro* assays were balanced at pH 7, which limited toxicity that might be present in areas of the gut with lower pH and higher fermentative metabolic activity (32). Further research could disentangle the effects of butyrate on *C. difficile* physiology at unique pH levels (32).

Several studies have observed the presence of butyrogenic pathways in *C. difficile*, which may use butyrate as a terminal electron acceptor in the absence of other options, such as Stickland amino acids (55-57). Due to unique toxicity patterns among the types of media used in our study as well as a previous study, we hypothesize that butyrate may have unique effects on *C. difficile* depending on which metabolic pathways are in use at the time of exposure (20, 58). This may be due in part to pressure from a build-up of downstream metabolic by-products as has been observed in *E. coli’s* response to high concentrations of acetate and formate (59). Further work is required to characterize the specific effects of SCFAs on *C. difficile* physiology and potential impacts on virulence (53). In summary, if butyrate is one of the mediators of increased colonization resistance in the inulin-treated communities, our data suggest that it is acting in concert with other unknown mechanisms.

Understanding the establishment of microbes in a new environment is essential as we seek to develop defined consortia to treat microbiota-related gut conditions, such as*C. difficile* infection. Some of the limited success of certain consortia may be due in part to low seeding densities as well as the inadequacy of smaller consortia to cover the appropriate niche spaces. Keeping these ecological dynamics in mind will assist in creating reliable treatments with broader efficacy across a population.

## 4. Materials and Methods

### 4.1. Stool collection

This study was approved by the University of Michigan’s Institutional Review Board (IRB: HUM00141992). We recruited adults (age > 18) and who had no history of gastrointestinal disease, including IBD, IBS, Crohn’s Disease and cancer. Individuals also had no history of antibiotic use or intestinal infection (bacterial or viral) in the previous six months. Exclusion criteria included immunocompromised status and immunosuppressant use. Following informed consent, we provided enrolled subjects with a commode specimen collection hat, and conical tubes, and instructions to freeze the stool sample, if delivery and sample collection exceeded the given timeframe. Upon receipt of each sample, we compensated subjects, and stored the sample at -80°C until use. For the experiments in this manuscript, we recruited two male individuals, aged 29 and 32.

### 4.2. Bioreactor set-up and operation

Through a collaboration with Robert Britton (Baylor University, Houston, TX), we received three-dimensional designs for bioreactor strips, each containing six reactors (14). With these designs, Protolabs, Inc. (Maple Plain, MN) used stereolithography to create each bioreactor strip from a thermostable resin (Somos WaterShed XC 11122). The operation of these bioreactor arrays has been previously described (14). Briefly, we filled reactors with 15 mL of bioreactor media (BRM) prepared as previously described, except that we sterilized bovine bile (Sigma, St. Louis, MO) by filtering at 0.22 μm (14). Once we established continuous flow, multichannel Watson Marlow peristaltic pumps (Falmouth, UK) individually maintained media flow to each reactor (1 rpm, 0.89 mm bore tubing) at a rate of 0.13 mL/hr.

### 4.3. Bioreactor dilution experiment

To prepare the fecal inoculum, we suspended fecal content from Subject A in sterile, pre-reduced phosphate-buffered saline (PBS; Thermo Fisher) at a ratio of 1:2. Feces were serially diluted by 10-fold to 10^−3^, 10^−5^, 10^−7^, and 10^−9^ in sterile, pre-reduced PBS and established in reactors in sextuplicate (N = 4 for 10^−3^). After 24 hrs of static culture, we initiated continuous flow and allowed to grow for 6 days before challenging with vegetative *Clostridioides difficile* str. 630 (Fig. 1A). Immediately before *C. difficile* challenge, we screened all reactors for possible contamination by plating on cycloserine-cefoxitin-fructose agar containing 0.1% taurocholate (TCCFA), which we made as previously described (60). To prepare *C. difficile* for challenge, we streaked spores onto agar plates containing taurocholate for 1 day at 37°C under anaerobic conditions. After incubation, we inoculated a *C. difficile* colony into 10 mL of sterile, pre-reduced BRM. At approximately 16 hrs of incubation, we back-diluted 1 mL of the culture in BRM by 10-fold and monitored to ensure *C. difficile* was in a log-phase of growth. Upon reaching OD 0.1, we again back-diluted the *C. difficile* culture in BRM and then inoculated into each reactor. We took 1 mL samples from the reactors at days 0 (i.e., the start of flow), 2, 3, 6, 7, 9 and 10. We immediately pelleted cells, and transferred the supernatant to be stored separately at -80°C. To assess *C. difficile* colonization, we enumerated colony-forming units (CFU) by serial dilution and plating on TCCFA.

### 4.4. Bioreactor carbohydrate experiment

To prepare the fecal inoculum, we suspended fecal content from Subject B in PBS as described above and then serially diluted to 10^−3^, 10^−4^, 10^−5^ and 10^−6^. We established fecal dilutions into the bioreactors in triplicate. After 24 hrs of static culture, we established continuous flow using BRM (Fig. 4A). After 48 hrs of continuous flow, we switched source media for the reactors in each dilution group to BRM containing 10-fold higher inulin (Swanson, Fargo, ND) than standard BRM, increasing inulin concentrations from the 0.02% present in basal media to 0.2%. We held conditions for 5 days to give reactor communities time to establish before challenging with *C. difficile* on day 7 as described above. We took 1 mL samples at days 0, 2, 5, 6, 7, 8, 10 and 11. During sampling, we pelleted cells and transferred the supernatant to store at -80°C. We then resuspended the pellet in RNAprotect Bacteria Reagent (Qiagen, Germantown, MD). We measured pH on day 10 of the experiment using MColorpHast pH test strips (Sigma).

### 4.5. DNA extraction and 16S rRNA-gene sequencing

We followed a detailed protocol for DNA extraction and Illumina MiSeq sequencing as described in previous publications with modifications (60). For the dilution bioreactor experiment, we pelleted cells and froze them at -80°C. In preparation for sequencing, cells were bead beaten in molecular grade water using 0.1 mm silica beads for 2 minutes. We then submitted cell extracts to the University of Michigan Microbiome Core for sequencing. For the carbohydrate bioreactor experiment, we pelleted cells and resuspended in Qiagen RNAProtect Bacteria Reagent before storing at -80°C. In preparation for sequencing, we submitted samples to the University of Michigan Microbiome Core for extraction using the Qiagen MagAttract PowerMicrobiome DNA/RNA Isolation Kit. For both experiments, we randomized samples into each extraction plate. To amplify the DNA, we used barcoded dual-index primers specific to the V4 region of the 16S rRNA-gene, and ran negative and positive controls in each sequencing plate (61). We prepared and sequenced libraries using the 500-cycle MiSeq V2 reagent kit (Illumina, San Diego, CA). Raw FASTQ files were deposited in the Sequence Read Archive (SRA) database (BioProject Accession: PRJNA837590).

### 4.6. Data processing and microbiota analysis

We performed 16S rRNA-gene sequencing as previously described using the V4 variable region and analyzed using mothur (62). Detailed methods, processed read data and the data analysis are described on GitHub (https://github.com/mschnizlein/cdiff_foundereffects). Briefly, after initial steps, such as assembly and quality control, we aligned contigs to the Silva v. 138 16S rRNA database (63). We removed chimeras using UCHIME and excluded samples with less than 1000 sequences (64). We binned contigs into operational taxonomic units (OTUs) by 97% percent similarity using Opticlust and used the Silva rRNA sequence database to classify those sequences (63, 65). Alpha and beta diversity metrics were calculated from unfiltered OTU samples. After subsampling to 2000 sequences, we used Dirichlet Multinomial Modelling (DMM) to identify bacterial enterotypes based on genus-level classification and then used LEfSe to identify taxa that significantly associate with each of these community types (66, 67). We performed all statistical analyses in R (v. 4.1.1) using the following packages: ggplot2, reshape2, plyr, tidyverse, ComplexHeatmap, and scales (68-73).

### 4.7. 16S rRNA-gene qPCR

Using dilutions of *Escherichia coli* ECOR2 genomic DNA as standards, we performed qPCR using PrimeTime gene expression master mix (IDT, Coralville, IA) and a set of broad-range 16S rRNA gene primers on a Thermo Fisher QuantStudio 3 (74). The DNA samples, standards, and negative controls were all amplified in triplicate. The qPCR reaction conditions were as follows: 95°C for 3 min, followed by 40 cycles of two-step amplification at 95°C for 15 s and 60°C for 60 s. The quantification cycle (Cq) values for each reaction were determined by using the Thermo Fisher Cloud software, and sample DNA concentrations were determined by comparing Cq values to the standards in each plate.

### 4.8. *C. difficile* growth curves

We isolated *C. difficile* strain 630g from a spore stock by growing overnight on BHI agar (BD) supplemented with 0.01% L-cysteine hydrochloride monohydrate (BHI; Sigma) and 0.1% taurocholate (Sigma). Growth curves were conducted with two biological replicates grown from two unique colonies. After growing overnight in BHI, we back-diluted cultures in fresh BHI with the overnight sample, and optical density was monitored to ensure cultures were in log-phase growth. Prior to the growth assay, we pelleted the culture, and then resuspended it in fresh 2x concentration BHI. We mixed this bacterial suspension into sodium butyrate solutions buffered at pH 7 with PBS ranging from 160mM to 2.5mM (2x final concentrations). The cultures were then placed in a 96-well plate optical density reader (Tecan, Switzerland) and monitored for 24 hours. All conditions were run with three technical replicates. Optical density measurements at 600 nm were automatically taken every 15 min, with 60 s of shaking immediately prior to measurement. We repeated this protocol in a follow-up experiment but substituted BHI for BRM in all steps following *C. difficile* colony isolation.

### 4.9. Short-chain fatty acid analysis

100 uL of fecal supernatants were filtered using 0.22 micron 96-well filter plates and stored at -20°C until analysis. We transferred the filtrate to 1.5 mL screw cap vials with 100 uL inserts for high performance liquid chromatography (HPLC) analysis and then randomized them. We quantified acetate, propionate, and butyrate concentrations using a refractive index detector as part of a Shimadzu HPLC system (Shimadzu Scientific Instruments, Columbia, MD) as previously described (75). Briefly, we used a 0.01 NH_2_SO_4_ mobile phase in filtered, Milli-Q water through an Aminex HPX87H column (Bio-Rad Laboratories, Hercules, CA). Sample areas under the curve were compared to volatile fatty acid standards with concentrations of 40, 20, 10, 5, 2.5, 1, 0.5, 0.25, and 0.1 mM. Through blinded curation, we assessed baseline and peak quality and excluded poor quality data if necessary.

---

**Figure S1: Individualized establishment dynamics of diluted microbial communities. A)** Observed species (SOBS) from all timepoints are plotted by community dilution. **B)** 16S rRNA-gene copies on days 0 and 6 are plotted by dilution. **C-F)** Relative abundance of 16S rRNA gene sequences is plotted for each individual reactor in the communities diluted **C)** 10^−3^, **D)** 10^−5^, **E)** 10^−7^, and **F)** 10^−9^ (Bacteroidota = green, Firmicutes = blue/purple, Actinobacteriota = orange, Proteobacteriota = red and Verrucomicrobiota = yellow. Other phyla, low abundance taxa and unclassified bacteria are colored as grey).

**Figure S2: Intra-reactor community variability by dilution**. Intra-reactor β-diversity was compared across samples within each reactor for the four community dilution groups using the **A)** Bray-Curtis Dissimilarity Index and the **B)** Jaccard Dissimilarity Index (statistical analysis: one-way ANOVA).

**Figure S3: Associations between *C. difficile* and resident microbes. A)** Observed species (SOBS) in each community dilution following *C. difficile* challenge. *C. difficile* abundance compared to the SOBS on **B)** day 6 and **C)** day 7 (statistical analysis: linear regression). LEfSe analysis comparing bacterial taxa in community types 1 & 3 with those in type 2. Certain taxa were more abundant in **D)** types 1 & 3 and others in **E)** type 2.

**Figure S4: Effects of inulin on microbial community function. A)** Bray-Curtis Dissimilarity index of communities in the 0.2% inulin treated reactor groups colored by dilution. Each point represents the distance of each reactor community relative to its baseline at day 0 (statistical analysis: Wilcoxon Rank Sum Test comparing distance to baseline at day 2 and day 5; for 10^−3^, *p* = 0.016; for 10^−4^, 10^−5^ and 10^−6^, *p* = not significant). **B)** pH of each reactor by dilution group at day 10 (statistical test: Kruskal-Wallis test). **C)** Butyrate concentrations at day 2 (pre-inulin shift) compared to concentrations at day 7 (statistical analysis: linear regression). **D)** Butyrate concentrations at day 7 compared ton *C. difficile* colonization on day 11. *C. difficile* growth curves in assessing butyrate toxicity in **E)** BHI and **F)** BRM, each buffered at pH 7.

## Acknowledgements

We would like to thank the Microbial Systems and Molecular Biology Core at the University of Michigan for their work performing the DNA extractions and 16S rRNA-gene sequencing. Many thanks go to Deanna Montgomery for critical review of this manuscript. We would also like to thank Thomas Schmidt and Kwi Kim for their guidance performing the HPLC short chain fatty acid analysis. Finally, we thank members of the Robert Britton lab for assistance as we got the bioreactor model running in the lab. This work was funded by the National Institutes for Allergy and Infectious Disease (U01-AI124255) and for Diabetes and Digestive and Kidney Diseases (T32-DK094775).

## Contributions

M.K.S.: Conceptualization, data curation, formal analysis, investigation, methodology, project administration, visualization, writing (original draft), writing (review & editing) A.K.S.: investigation, methodology, writing (review & editing)

M.J.G.: investigation, formal analysis, visualization, writing (review & editing)

S.J.E.: investigation, formal analysis, writing (review & editing)

V.B.Y.: Conceptualization, funding acquisition, project administration, resources, supervision, writing (original draft), writing (review & editing)

