## Supplementary figures and images for "Differences in Gut Microbiota Assembly Alter Its Ability to Metabolize Dietary Polysaccharides and Resist *Clostridioides difficile* Colonization"

### Supplemental Figure 1

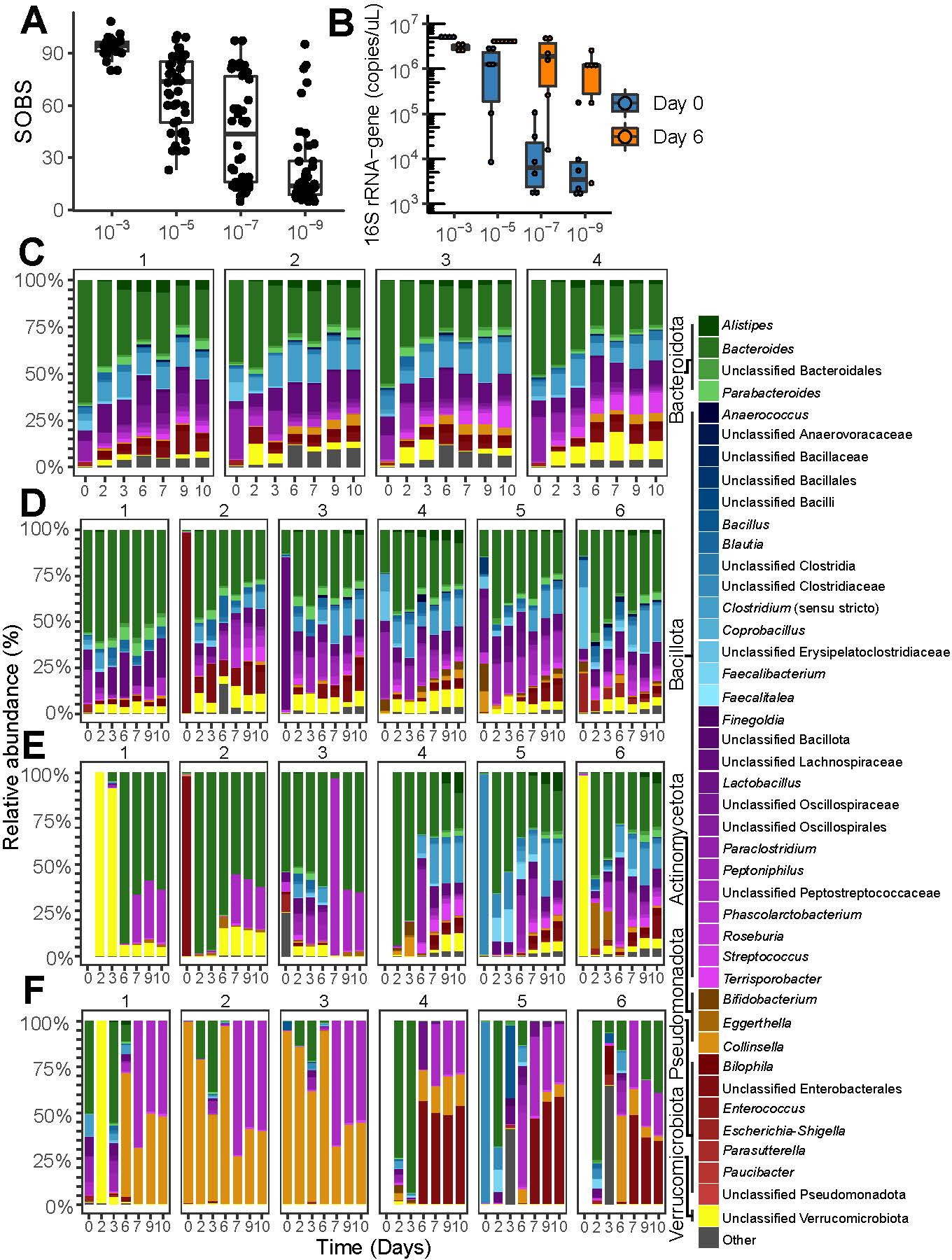

### Supplemental Figure 2

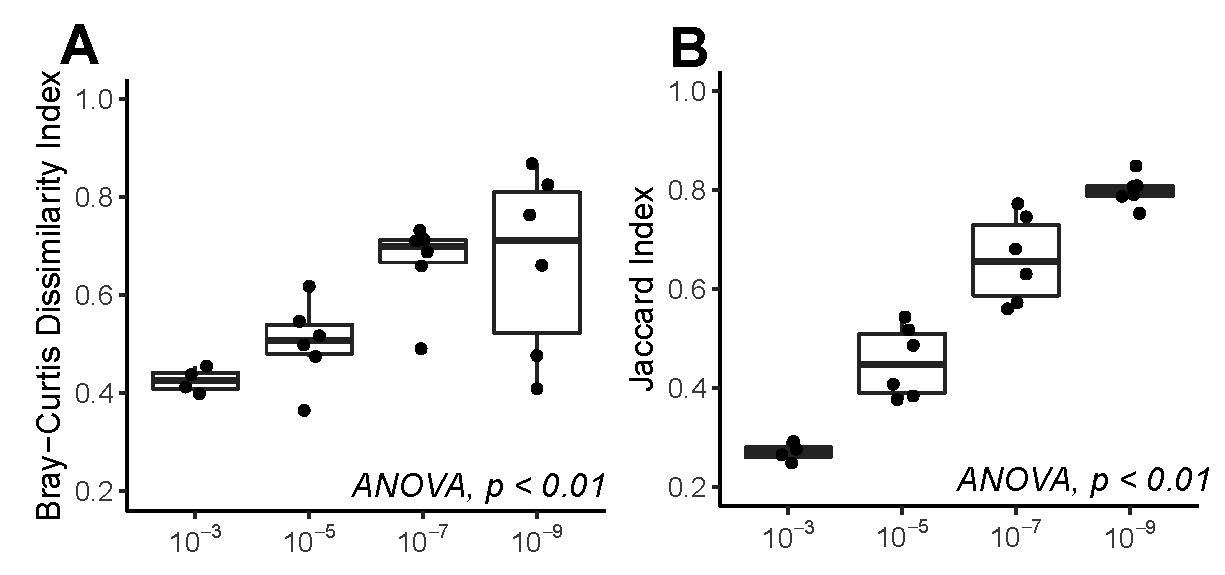

### Supplemental Figure 3

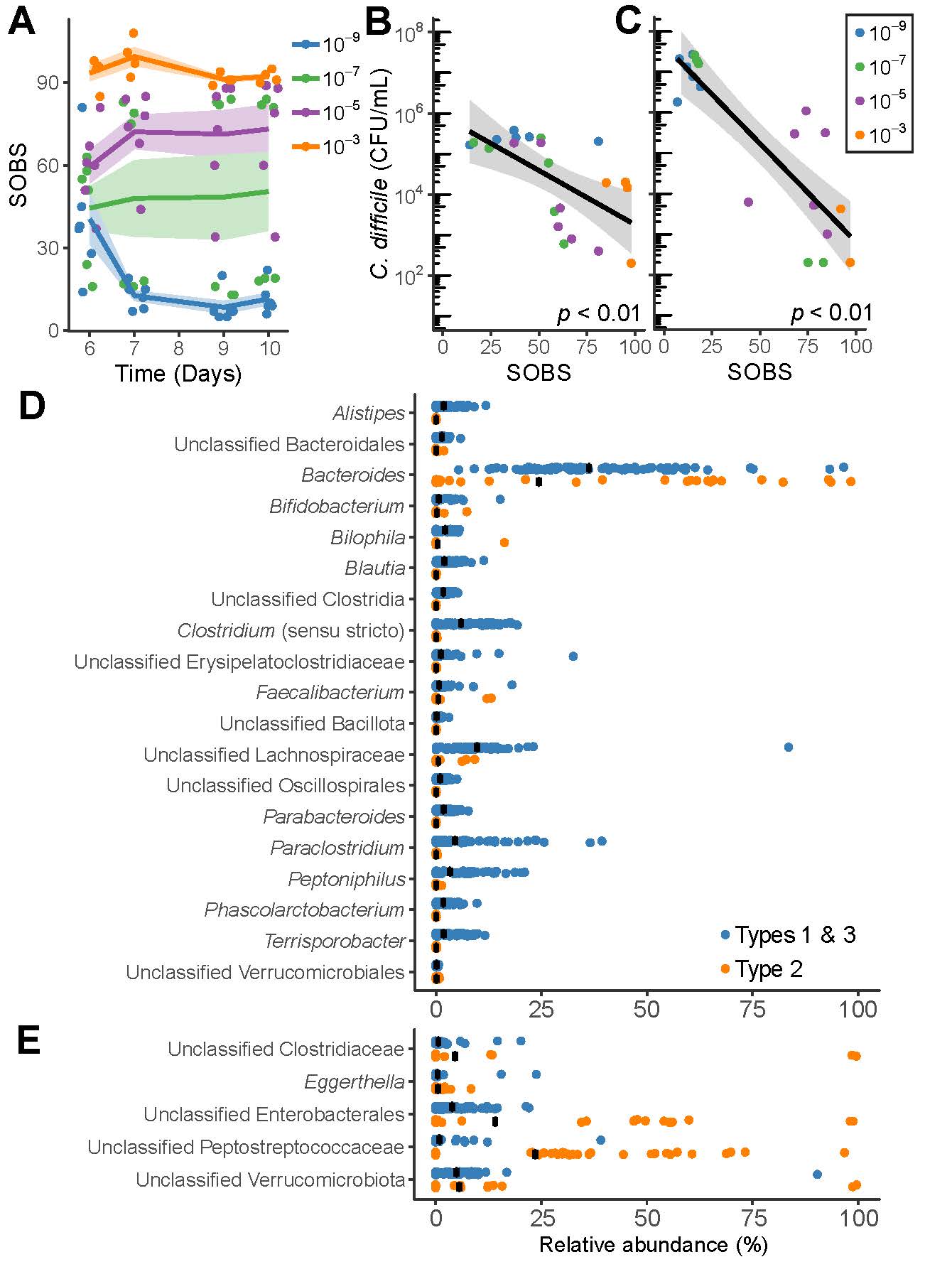

### Supplemental Figure 4

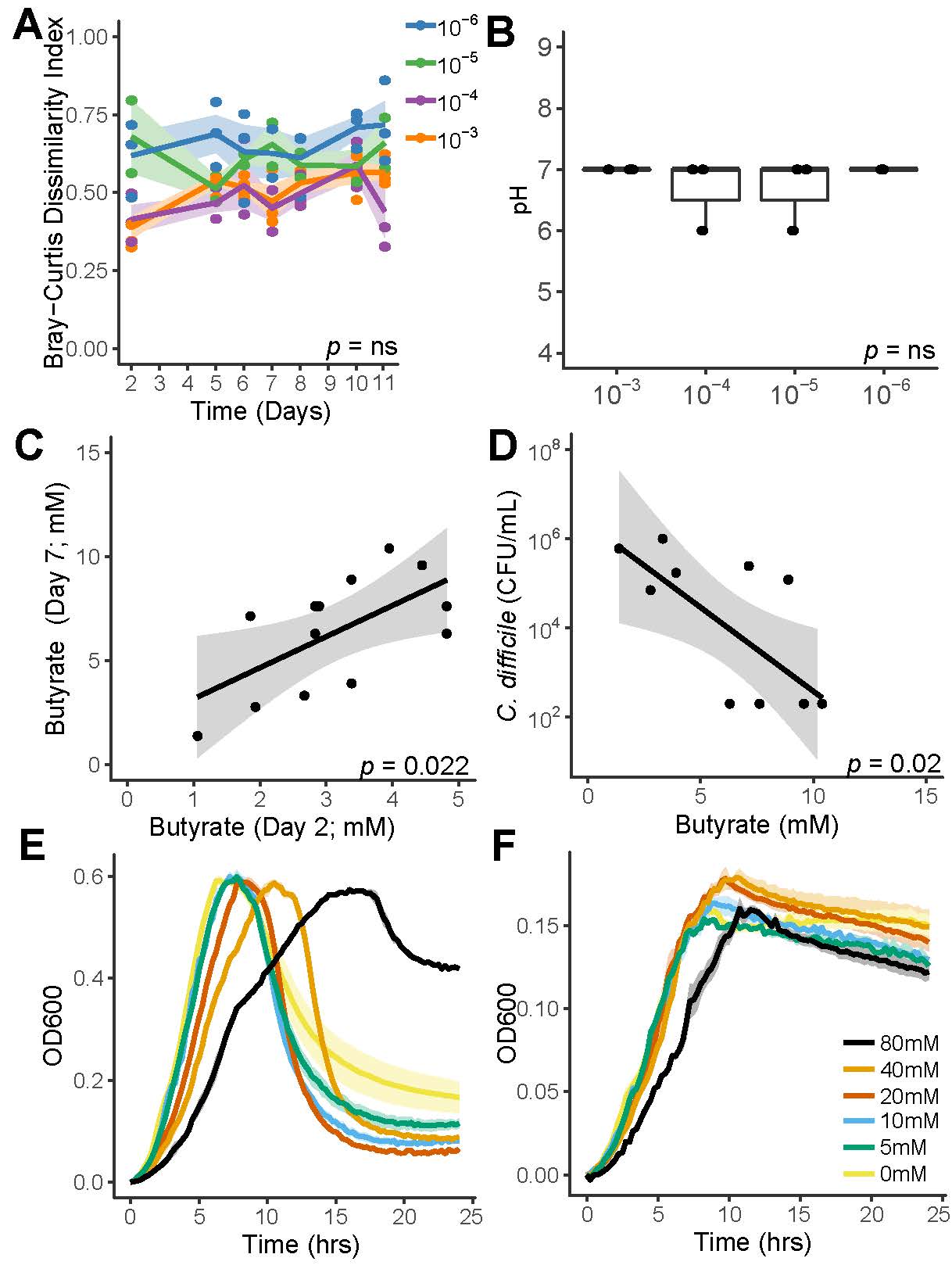
